# Recognition and deubiquitination of free 40S for translational reset by Otu2

**DOI:** 10.1101/2021.11.17.468975

**Authors:** Ken Ikeuchi, Nives Ivic, Jingdong Cheng, Robert Buschauer, Yoshitaka Matsuo, Thomas Fröhlich, Otto Berninghausen, Toshifumi Inada, Thomas Becker, Roland Beckmann

## Abstract

In actively translating 80S ribosomes the ribosomal protein eS7 of the 40S subunit is monoubiquitinated by the E3 ligase Not4^1,2^ and deubiquitinated by the deubiquitination enzyme Otu2 upon ribosomal subunit recycling^3^. Despite its importance for general efficiency of translation the exact role and structural basis for this specific translational reset are only poorly understood. Here we present biochemical and structural data showing that Otu2 can engage the recycled 40S subunit together with the recycling factors ABCE1 and Tma64 immediately after 60S dissociation for mRNA recycling, and that it dissociates before 48S initiation complex formation. A combined structural analysis of Otu2 and Otu2-40S complexes by X-ray crystallography, AlphaFold2 prediction^4^ and cryo-EM revealed how Otu2 can specifically be recruited to the 40S, but not to the 80S ribosome, for removal of the eS7-bound ubiquitin moiety. Here, interactions of the largely helical N-terminal domain of Otu2 to sites that are masked and therefore inaccessible in the 80S ribosome are of crucial importance. Collectively, we provide the structural basis for the Otu2 driven deubiquitination step providing a first mechanistic understanding of this translational reset step during ribosome recycling/(re)initiation.

## Introduction

The addition of ubiquitin to lysine residues of a protein or to ubiquitin itself is a very common post-translational modification. It creates multi-faceted cellular signals, known as the ‘ubiquitin code’ that can lead to a multitude of possible cellular consequences such as protein degradation, signaling and trafficking, epigenetic regulation, cell cycle control and many more^5^. Commonly, ubiquitin is added via sequential action of three enzyme classes, E1 ubiquitin-activating enzymes, E2 ubiquitin-conjugating enzymes and E3 ubiquitin ligases. The effects of ubiquitination are counteracted by deubiquitinating enzymes (DUBs), that are able to remove (poly)ubiquitin chains^6–7^.

Recently, important roles of ubiquitination and deubiquitination have emerged in the context of eukaryotic translation. For example, small subunit proteins uS3, uS10 and eS10 are ubiquitinated following ribosome collision after prolonged stalling of translation^8 9 10 11 12 13 14^. This can trigger mRNA surveillance and ribosome quality control (RQC) pathways, during which stalled ribosomes are dissociated into subunits and the mRNA as well as the arrest peptide are degraded^15 16 17 18 19 20^. uS3 and uS5 are also ubiquitinated under several types of translational stress^21 22 23 24^ and sequential ubiquitination of uS3 can promote the 18S non-functional ribosomal RNA decay (NRD) pathway^25^. An important target for general translation coordination is eS7, that in yeast is monoubiquitinated by the E3 ligase Not4 ^1 2 9^, a component of the Ccr4-Not complex and master regulator of gene expression^26^. eS7 ubiquitination by Not4 occurs constitutively after canonical translation initiation and it was demonstrated to be required for mRNA homeostasis by serving as a prerequisite of Ccr4-Not’s role in degradation of non-optimal mRNAs^1–27^.

Monoubiquitination of eS7 is also responsible for translational regulation under ER stress condition^28^. Moreover, in yeast, eS7 can be polyubiquitinated by the E3 ligase Hel2 to trigger a bypass quality control pathway^9^. A recent study revealed, that eS7 poly- and monoubiquitination is antagonized by deubiquitinating enzymes Ubp3 (USP10 in human) and Otu2^3,28^. While Ubp3 preserves the eS7 monoubiquitination, Otu2 was shown to specifically remove the monoubiquitin attached to K83 of eS7^3^. This step exclusively occurs on 40S ribosomal subunits and was suggested to happen during recycling of mRNA and tRNA from 40S after translation termination^3^.

Yet, several questions remain open. First, given that productively translating ribosomes require monoubiquitination of eS7, Otu2 activity must not occur on 80S ribosomes but remain limited to free 40S subunits. However, considering that eS7 is equally accessible in 40S and 80S ribosomes, it is not clear at which stage of the translation cycle Otu2 is active and how 40S specificity is achieved.

## Results

### Otu2 association with 40S ribosome recycling and translation initiation complexes

To explore the role of ribosome-bound Otu2 and Ubp3, and to identify additional factors associated with the respective complexes, we performed a quantitative mass spectrometry-based analysis. To that end, we affinity purified Otu2-FTpA (FLAG-TEV-proteinA) and Ubp3-FTpA associated complexes from yeast cells (Fig. 1a, b) and analyzed them by mass spectrometry (Fig.1c, Supplementary Table 1). In the Ubp3-bound fraction, primarily its co-factor Bre5 (G3BP1 in human)^29^ as well as large ribosomal subunit proteins were enriched, in agreement with a suggested function of Ubp3 on 80S ribosomes^3 28 30 31 32 33^. In addition, we found factors involved in ER to Golgi trafficking which is in agreement with a known function of Ubp3 independent of translation^29,34^. In the Otu2-associated fraction, however, we found enrichment of small ribosomal subunit proteins and of factors involved in translation termination, ribosome and mRNA recycling (ABCE1; Rli1 in yeast, release factors, Tma64; eIF2D in human) as well as proteins involved in small subunit biogenesis (Fig. 1c, Supplementary Table 1). In addition, translation initiation factors (subunits of the eIF2 and eIF3 complexes) were enriched in the wash fraction after micrococcal S7 nuclease treatment, that contained mRNA-mediated or less stably associated factors (Fig. 1d). Independent of its role in translation, Otu2 is likely to play an additional role in 40S biogenesis since this was shown for the human Otu2 homolog, OTUD6B^35^. Although a functional role of eS7 (de)ubiquitination during pre-40S formation in yeast has not yet been described, the enrichment of biogenesis factors in Otu2 complexes supports this idea. The observation of strong enrichment of 40S ribosomal proteins clearly indicates that in contrast to 80S associating Ubp3, Otu2 binds with high preference to 40S subunits. Based on the enrichment of the recycling factors ABCE1 and Tma64, Otu2 association occurs early during recycling, immediately after ABCE1-induced dissociation of the 60S subunit^36^ and during dissociation of the mRNA facilitated by factors like Tma64^37 38 39^. This is consistent with earlier biochemical findings that the impairment of Otu2 activity results in a reduced efficiency of mRNA recycling^3^. The presence of Otu2 on the recycled 40S can last into the phase of 43S pre-initiation complex formation as indicated by the presence of eIF2 and eIF3 subunits. This may hint at an additional function of Otu2 after mRNA recycling during early phases of (re-)initiation. However, the lack of enrichment of any 48S factors such as eIF4A, eIF4G or eIF4E suggests that Otu2 dissociates from the 40S subunit before 48S formation and mRNA scanning.

**Fig. 1.**
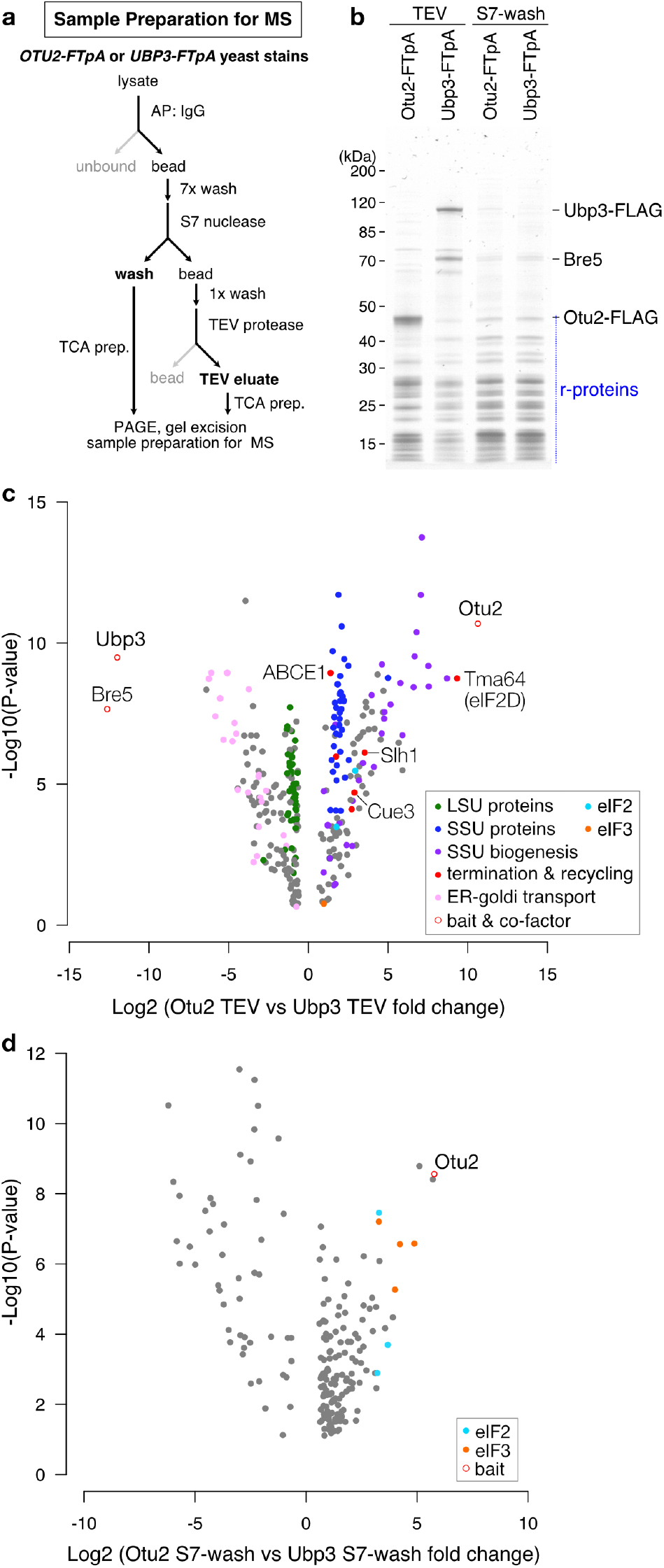
Quantitative mass spectrometry-based analysis of ribosome-bound Otu2 versus Ubp3. a, Scheme outlining the sample preparation for mass spectrometry. MS; mass spectrometry, AP; affinity purification, TCA; trichloroacetic acid, TEV; tobacco etch virus. b, Nu-PAGE gel of affinity purified fractions by Otu2-FTpA and Ubp3-FTpA. c, Volcano plot displaying enrichment of copurified proteins determined by mass spectrometric analysis for TEV elution, and d, wash samples after S7 nuclease treatment.

### Otu2 binds to 40S subunits for deubiquitination of monoubiquitinated eS7

Next, we further analyzed ribosome association and eS7 deubiquitination activity of Otu2 and catalytically inactive mutants in yeast cell lysates. Therefore, we expressed C-terminally 3x FLAG-tagged Otu2, otu2-C178S and otu2-H300A from plasmids under the endogenous promoter in yeast strains with *otu2*Δ or *otu2Δubp3Δ* background harboring shuffled hemagglutinin (HA)-tagged eS7A to monitor ubiquitination. Western Blot analysis of lysates subjected to sucrose density gradient fractionation confirmed that both active and inactive Otu2 are associated with 40S subunits and that deubiquitination of eS7 on 40S is dependent on the catalytic activity of Otu2 (Supplementary Fig. 1a, b). Moreover, catalytic inactivity is accompanied by a decrease of polysomes, emphasizing the proposed importance of Otu2 function for efficient translation (Supplementary Fig.1c). This is further supported by the fact, that the previously observed synthetic growth defect of *otu2*Δ*ubp3*Δ cells^3^ is even more pronounced in the presence of translation elongation inhibitors or at higher temperature (Supplementary Fig. 1d).

### Cryo-EM structure of the Otu2-40S complex

To gain mechanistic insights into the role of Otu2 in translation, we generated an Otu2-40S complex sample *in vitro* for cryo-EM analysis. To that end, we reconstituted an Otu2-eS7-Ub-40S complex using an *in vitro* ubiquitination system. Herein, purified 80S ribosomes were monoubiquitinated with purified Not4 E3 ligase, Ubc4 (E2), Uba1 (E1), ubiquitin and ATP to obtain 80S with monoubiquitinated eS7 (eS7-Ub-80S; Fig. 2a, b, Supplementary Fig. 2). These ribosomes were split into subunits (Fig. 2b) and purified eS7-Ub-40S were incubated with purified recombinant Otu2 to ensure efficient deubiquitination activity in this system (Fig. 2c). After observing complete eS7-de-ubiquitination by the purified wild-type (wt) Otu2 we used the eS7-Ub-40S for reconstitution with a 5x molar excess of purified catalytically inactive otu2-C178S to form stable complexes (Fig. 2d) and subjected them to cryo-EM.

**Fig. 2.**
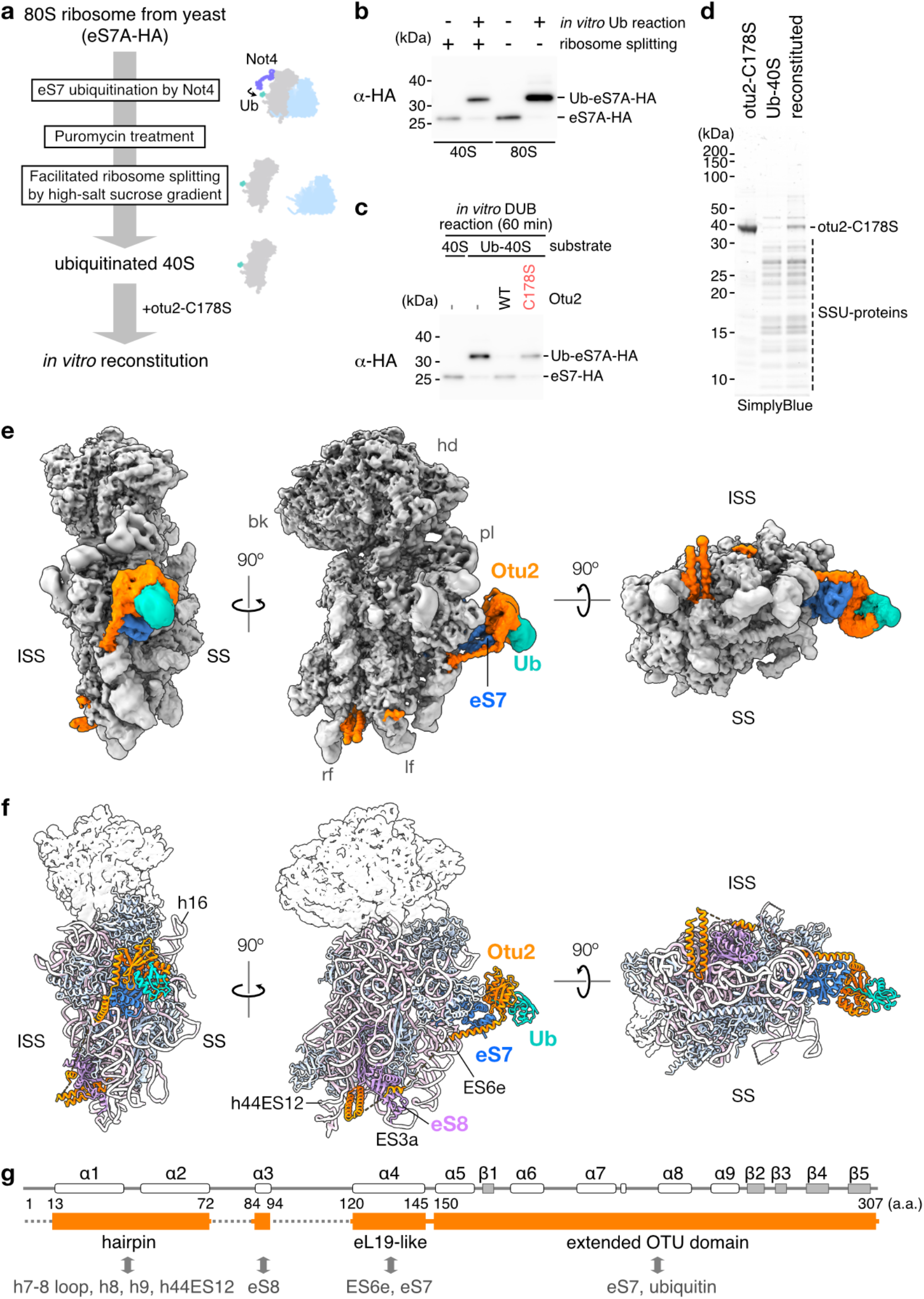
Cryo-EM structures of Otu2-40S complexes. a, Scheme outlining the *in vitro* generation of eS7A-monoubiquitinated 40S. b, a-HA Western Blot showing eS7A-ubiquitinated 40S after puromycin/high salt induced splitting of 80S ribosomes. c, a-HA Western Blot showing *in vitro* deubiquitination reactions with eS7A-monoubiquitinated 40S and wt Otu2 or otu2-C178S mutant protein. d, Nu-PAGE gel showing the Otu2-40S cryo-EM sample *in vitro* reconstituted from purified eS7A-monoubiquitinated 40S subunits and otu2-C178S mutant protein. e, Cryo-EM structure of the *in vitro* reconstituted Otu2-40S complex. Non-ribosomal extra densities for the Otu2 OTU domain (orange) and ubiquitin (light sea green) are visible close to eS7 (blue). Additional extra density for the Otu2 N-terminus is visible close to the 40S foot structure. The cryo-EM map was low-pass filtered according to local resolution. ISS; intersubunit surface, SS; solvent surface, hd; head, bk; beak, pl; platform, rf; right foot, lf; left foot. f, Molecular model for Otu2, monoubiquitinated eS7 and the 40S body. The 40S head is shown as transparent density. g, Schematic illustration of Otu2 secondary structure and domains. Structurally visible parts are highlighted as orange boxes.

A 3D reconstruction of the Otu2-40S complex was obtained after 3D classification and multi-body refinement^40^ (Supplementary Figs. 3, 4a and 4b, Table 1>). After low-pass filtering according to local resolution (ranging between 2.8 Å for the ribosomal core and 11 Å for flexible elements), the structure revealed extra density for the OTU domain (167-305) and ubiquitin adjacent to eS7 (Fig. 2e). Similar to OTUD1 and OTUD3, the OTU domain is extended on the N-terminal side by an a-helix (a5; 150-166, Supplementary Figs. 5, 6) that contacts eS7. In addition, we observed parts of the Otu2-specific N-terminus (1-149) which winds along the lower part of the 40S body on the intersubunit side (ISS) connecting the ‘foot’ structure (formed by the lower parts of rRNA helix h44 that extends into expansion segment ES12) with the OTU domain (Fig. 2e).

We were able to build a partial model for the four a-helices (α1-α4) based on a model generated by AlphaFold v2.0 (AF2; Fig. 2f, g, Supplementary Fig. 4d, e and 5a-c)^4^: a1 and α2 form a hairpin that is inserted into a deep rRNA pocket formed by residues of h7-h8 loop, h9/ES3a and the lower end of h44/ES12 (Fig. 3a, Supplementary Fig. 4d). The hairpin is flexibly linked to the short a3 helix of Otu2 (also predicted by AF2) that binds on the surface of eS8. A longer flexible linker connects α3 with the long α4, that inserts into the major groove of the rRNA expansion segment ES6e and projects further towards eS7, where it connects to the OTU domain of Otu2 (Fig. 3b, Supplementary fig. 4e).

**Fig. 3.**
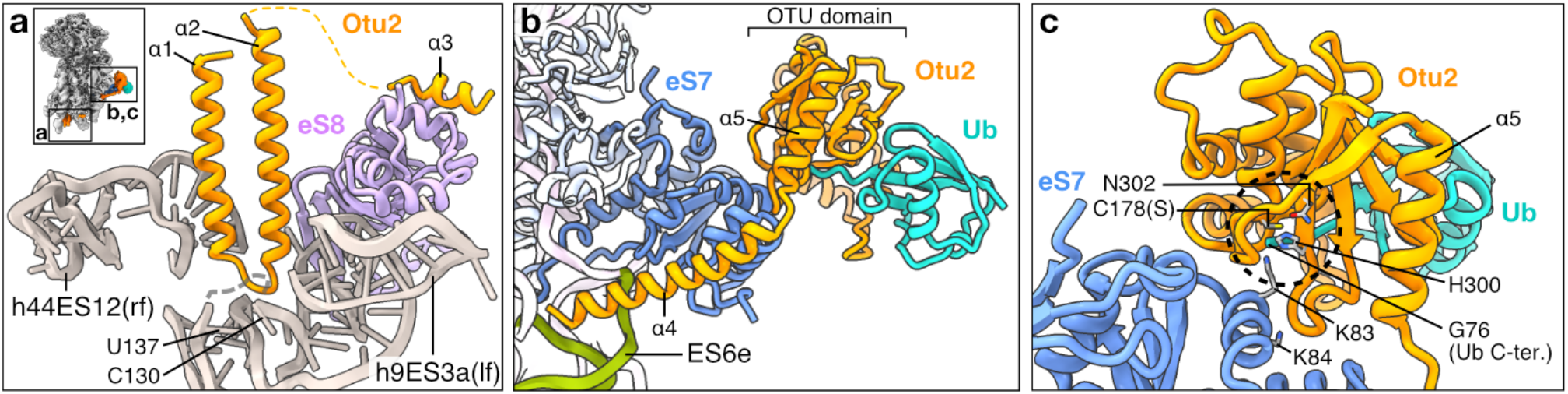
Molecular model of the Otu2-40S complex. a, View focusing on the N-terminal hairpin (α1-2) bound to h44 and α3 bound to eS8. rf; right foot, lf; left foot. b, View focusing on the Otu2 α4 and OTU domain bound to monoubiquitinated eS7. c, Detailed view focusing on the active site of Otu2 OTU domain targeting the ubiquitinated-eS7-K83 residue. The dotted circle shows the active protease site with the catalytic triad (C178, H300 and N302) of Otu2 and its substrate, ubiquitinated eS7-K83. The minor ubiquitination site eS7-K84 is also shown. A thumbnail in (a) indicates the location of zoomed region within the Otu2-40S complex.

Since local resolution of the OTU domain was not sufficient to build a *de novo* molecular model, we determined the structure of the N-terminally extended Otu2-OTU domain comprising residues 150-307 by X-ray crystallography (Table 2, Supplementary Fig. 5d). This structure was very similar to the OTU domain structure predicted by AF2 (Supplementary Fig. 5e, f; RMSD of 0.367) and fitted well with only minor adjustments into our cryo-EM density. This enabled us to obtain an overall model of eS7-mono-Ub bound Otu2 based on the crystal structure of ubiquitin-bound Otu1 (Fig. 2c, Supplementary Figs. 4e, f) (PDB: 3c0r)^41^.

The structure of the OTU domain of Otu2 showed the typical OTU domain fold and is most similar to human OTUD1 and OTUD3 domains (Supplementary Fig. 5g-i and 6), both of which have the N-terminal a-helix of the catalytic domain (α5 in Otu2; Asp149-Lys165; also called the S1’ helix), that is suggested to contribute to the ubiquitin linkage specificity and/or substrate specificity^42^. Examination of the position of the Otu2 OTU domain in our cryo-EM map revealed that the conserved core of the domain forms numerous contacts to K83-ubiquitinated eS7 involving loops α8-α9 (Trp248) and ß3-ß4 (Ser294-Gly298) adjacent to the catalytic triad (Cys178, His300 and Asn302) (Fig. 2c, Supplementary Fig. 5c, g).

Additional eS7 contacts are formed by the region linking the S1’-helix (α5) with α4 (140-151) (Fig. 2b). Although the local resolution was low due to flexibility, we observed a clear density for ubiquitin bound to the Otu2 S1 site superimposing well with ubiquitin-bound *S. c.* Otu1^41^ (3c0r) (Supplementary Fig. 4f). The position of the Otu2-bound ubiquitin on the 40S subunit localizes its C-terminal Gly76 in close proximity to the modified K83 (but not K84) of eS7. This finding is consistent with a K83-linked eS7-ubiquitin substrate complex engaged by the catalytically inactive Otu2^3,9^ (Fig. 3c).

In order to confirm the physiological relevance of our *in vitro* reconstituted complex, we also analyzed endogenously formed complexes of catalytically inactive otu2-C178S and 40S ribosomes. These complexes were obtained by purification from 40S fractions of otu2-C178S-3xFLAG-overexpressing yeast cells and subjected to structural analysis by cryo-EM (Supplementary Figs. 3b, 7a and 7b). Despite overall higher heterogeneity within this sample, we observed otu2-C178S-bound 40S subunits with an essentially identical architecture as observed for the *in vitro* assembled complexes (Supplementary Fig. 7c, d), including the N-terminal α1-α2 hairpin (Supplementary Fig. 7d, e).

### The N-terminal region of Otu2 provides specificity for 40S binding

Our structures suggest that the N-terminal region of Otu2 comprising helices α1-α4 may provide 40S binding specificity, since its presence on the 40S would sterically clash with the 60S subunit in context of an 80S ribosome. Most intriguingly, α4 occupies the binding position of LSU protein eL19, that forms the functionally important intersubunit bridge eB12^43^ (Fig. 4a, b). In 80S ribosomes eL19 inserts with its long C-terminal α-helix into the major groove of expansion segment ES6e. α4 of Otu2 occupies a very similar position on ES6e, most likely involving basic amino acids including a patch of four consecutive lysines and arginines (K116-R119; BP1) (Fig. 4b, Supplementary Fig. 5c). In addition to α4 of Otu2, that would clash with the 60S subunit, also the α1-α2 helical hairpin and α3 may contribute to 40S specificity (Fig. 4c, d). In context of the 80S ribosome, Otu2 binding is impaired by the tip of 25S rRNA H101 (ES41) of 60S subunit that is also involved in intersubunit bridge formation with eS8 (bridge eB11). Moreover, binding of the α1-α2 hairpin induces a small translational movement of the lower h44 towards the left foot to form the binding pocket. This would not be possible on an 80S ribosome, since h44 is involved in formation of the bridge eB13 with eL24 which stabilizes the overall 80S conformation.

**Fig. 4.**
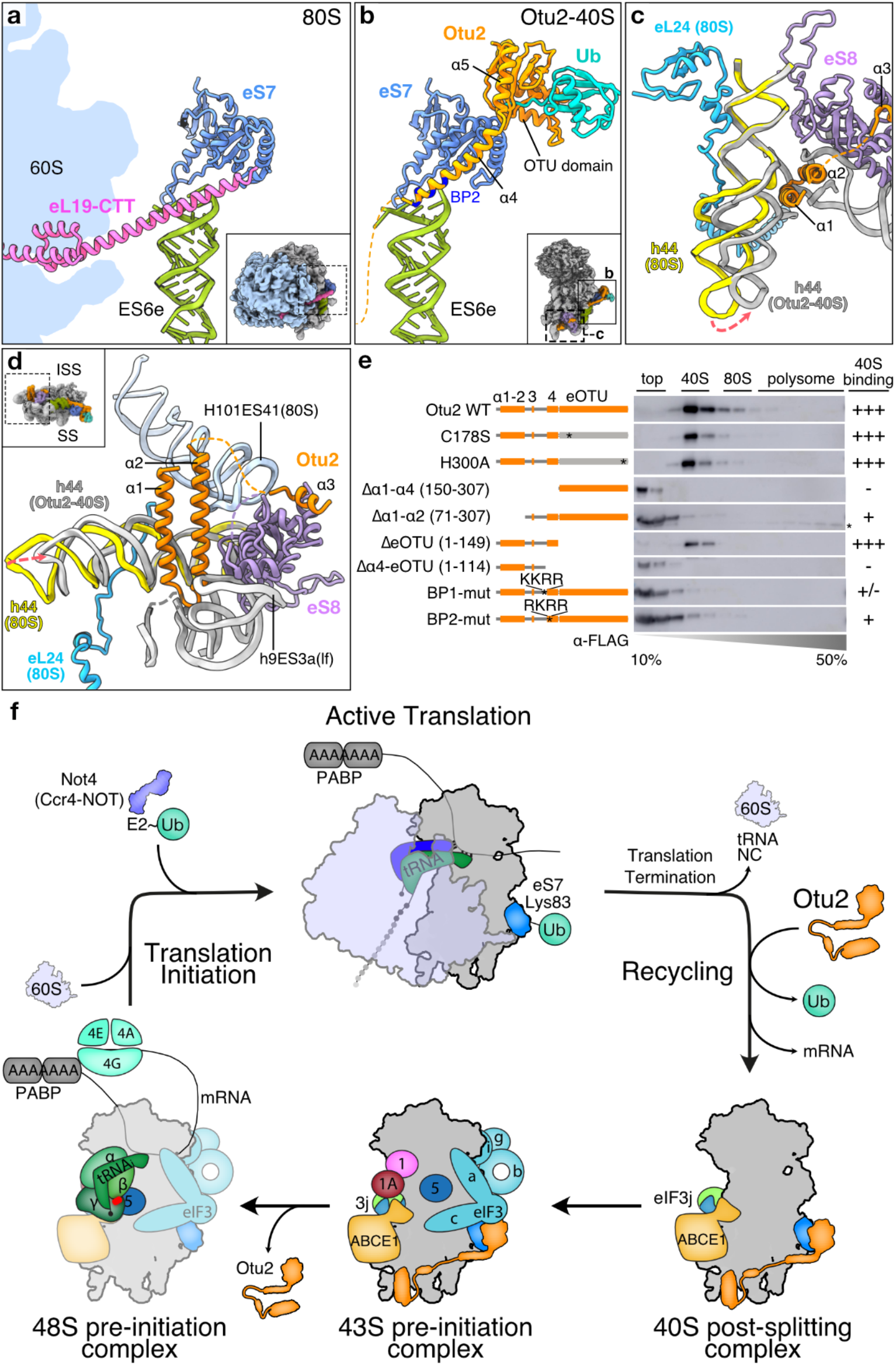
Comparison between the 40S subunit in context of 80S and bound to Otu2. a, View on intersubunit bridge eB12 formed by the C-terminal tail (CTT) of 60S protein eL19 and the 40S ES6e as present in the non-rotated state 80S (PDB:6snt). b, Same view as in a) on the Otu2-bound 40S subunit. Instead of eL19, α4 of Otu2 binds to ES6e major groove; residues contained in basic patches (BP2) are indicated. c and d, views focusing on the Otu2 α1-α2 hairpin and α3 located close to intersubunit bridges eB13 (formed by 60S protein eL24 and 40S h44) and eB11 (formed by 60S H101/ES41 and eS8 of the 40S subunit) in 80S non-rotated state. The hairpin binds to h44 opposite of the eL24 binding site. When bound to the hairpin, h44 moves towards the left foot (ES3a) thereby compacting the rRNA binding pocket for the hairpin. e, Western Blot analysis (α-FLAG) of fractions after sucrose density gradient centrifugation. Lysates were analyzed from an *otu2*Δ*ubp3*Δ yeast strain harboring a vector expressing the wt or indicated 3xFLAG-tagged Otu2 mutants. BP1-mut; K116A K117A R118A R119A, BP2-mut; R121A K123A R125A R129A. f, A scheme of the ubiquitination-deubiquitination cycle for yeast ribosomal protein eS7 during translation.

To test if these overlapping binding sites may indeed provide specificity for Otu2 recruitment to 40S subunits, we generated truncation mutants of Otu2 and checked their association with 40S or 80S ribosomes in sucrose density gradients (Fig. 4e). While the vast majority of the wt and catalytic triad mutant Otu2 (otu2-C178S and otu2-H300A) were found in the 40S fraction (Fig. 4e; Supplementary Fig. 1a), mutants lacking the entire N-terminus (Δα1-α4; 150-307) were not found in ribosomal fractions, indicating complete loss of 40S binding. Mutants without the α1-α2 hairpin (Δα1-α2; 71-307) but still containing α4 displayed some residual binding activity. This suggests that the OTU domain alone has insufficient affinity for stable 40S binding, and that recruitment of Otu2 to the 40S is mediated by its N-terminus with a major contribution by helix α4. In support of this, mutants lacking the extended OTU domain (ΔeOTU; 1-149, lacking α5) were still able to bind to 40S. However, shorter N-terminal fragments lacking α4 (Δα4-eOTU; 1-114) or carrying mutations in the basic patches close to or within α4 were not able to bind to 40S (BP1) or showed severely reduced binding (BP2) (Fig. 4b, e, Supplementary Figure 5c). Taken together, our data show that stable recruitment of Otu2 to the ribosome depends on its N-terminal α-helices, in particular helix α4. Since the interaction of the N-terminus with the 40S subunit is only possible in the absence of the 60S subunit, we conclude that these helices provide the specificity of Otu2 activity for 40S versus 80S ribosomes.

## Discussion/Conclusion

Collectively, our data indicate that for efficient translation Otu2 primarily acts during the 40S recycling and early (re-)initiation phase in agreement with its suggested role in mRNA recycling^3^ (Fig. 4f). This is plausible since our structure suggests that Otu2 can engage the 40S subunit as soon as 60S dissociation exposes the binding patches for the N-terminal domain of Otu2. Here, the major groove of ES6e of the 18S rRNA plays an important role. In the 80S ribosome it is occupied by eL19 of the 60S subunit, but on the free 40S subunit it can accommodate helix α4 of Otu2, which is necessary for efficient Otu2 binding. Ribosome binding of Otu2 should still be possible in the presence of recycling factors such as ABCE1 and Tma64/eIF2D, because none of the 40S-associating recycling and initiation factors occupy any of the Otu2 binding sites (Supplementary Fig. 8).

It is noteworthy that Otu2 remains bound to the 40S all the way into the 43S forming pre-initiation phase as indicated by the co-enrichment of eIF2 and eIF3 subunits. We therefore speculate that once bound to the recycled 40S, Otu2 may fulfill additional functions during translation initiation after ubiquitin removal from eS7 and mRNA recycling. Here the N-terminal α1-α2 helical hairpin is of particular interest since it is not strictly required for 40S binding but induces a conformational reorganization of the rRNA h44, thereby potentially contributing to mRNA recycling and 43S coordination by an allosteric mechanism.

Overall, our data provide the structural basis for the general translational reset step by deubiquitination of eS7 during recycling/(re)initiation. Yet, the precise interplay of Otu2 with the recycling/initiating 40S and the trigger for dissociation will be subject of future studies.

## Materials and Methods

### Yeast Strains and Plasmids

All yeast strains used in this study were derivatives of the budding yeast (*Saccharomyces cerevisiae*) W303-1a parental strain (see also Supplementary Table 2 and 3). For comparative mass spectrometric analysis, yeast strains expressing FLAG-TEV-ProteinA (FTpA)-tagged Otu2 and Ubp3 were generated by established yeast technique^44^. Gene deletion yeast strains were constructed by homologous recombination using amplified DNA fragments^45^. The coding sequence and the promoter of *OTU2* were cloned from W303-1a genomic DNA into the *p415-3xFLAG-CYC1t* vector. The Otu2 point mutants were generated by site-directed mutagenesis. N- or C-terminal Otu2 deletion mutants were created by PCR using specific oligonucleotides combinations. Otu2 expression plasmids were transformed into *otu2*Δ or *otu2*Δ*ubp3*Δ yeast strains using lithium acetate, then grown and selected on SDC -Leu medium plates. To obtain wild-type (wt) Otu2 and otu2-C178S, *OTU2* and *otu2-C178S* were cloned into a pGEX6P-1-His_6_-GST-3C expression vector. For recombinant Not4, Ubc4 and Uba1, pGEX6P-2-*NOT4-FLAG,* pGEX6P-1-*UBC4* vectors were used for expression in *E. coli* and the *p416GPDp-UBA1-FLAG* vector was used for expression in a yeast *leu1Δ* strain. To generate eS7A-HA-tagged 80S ribosomes, *otu2Δrps7aΔrps7bΔ* yeast strains harboring a eS7A-HA expressing plasmid were used as described previously^9^.

### *Ex vivo* pull-out of Otu2-40S complexes

Yeast cells expressing C-terminally 3xFLAG-tagged otu2-C178S were cultivated in 2 l SDC -Leu liquid medium, grown at 30°C until an OD_600_ of 1.0, harvested by centrifugation and quickly frozen in liquid nitrogen. The frozen yeast cell pellet was powdered in liquid nitrogen by grinding using a mortar and a pestle. Resulting yeast cell powder was resuspended in ice-chilled lysis buffer-DG (20 mM HEPES-KOH pH 7.5, 100 mM KOAc, 2 mM Mg(OAc)_2_, 1 mM dithiothreitol (DTT), 1 mM phenylmethylsulfonyl fluoride (PMSF), 1 tablet per 10 mL of cOmplete tablets EDTA-free (Roche)), spun down in a table top centrifuge to remove the rough cell debris, and further centrifuged at 40,000 *g* in a SS-34 rotor to remove debris. The yeast lysate was loaded on a 10-50% sucrose density gradient in SW32 tubes, and ultracentrifuged at 35,000 *g* for 16 h. The buffer of the harvested 40S fraction was changed to buffer-IP (20 mM HEPES-KOH pH 7.5, 100 mM KOAc, 10 mM Mg(OAc)_2_, 125 mM sucrose, 1 mM DTT, 0.01% NP-40) by using Amicon-100K, then anti-FLAG M2 magnetic agarose was added to the 40S fraction and incubated for 1 h on a nutator at 4°C. After washing the beads 5 times with buffer-IP, the beads were incubated with 400 μL buffer-IP containing 250 ng/μL 3xFLAG peptide for 1 h at 4°C to elute. The ribosomal complexes were purified by ultracentrifugation through a sucrose cushion in a TLA120.2 rotor at 355,000 *g* for 1 h and dissolving the ribosomal pellet with 30 μL buffer-EM (20 mM HEPES-KOH pH 7.6, 50 mM KOAc, 5 mM Mg(OAc)_2_, 50 mM sucrose, 1 mM DTT, 0.05% Nikkol). Purified samples were analyzed using SDS-PAGE and cryo-EM grids were prepared.

### *In vitro* reconstitution of Otu2-40S complexes

For the preparation of ubiquitinated 40S ribosomal subunits, the yeast eS7A-HA-shuffled strain was cultivated in 2 l of YPD and harvested at an OD600 of 1.5. Yeast crude ribosomes were purified from the yeast lysate by pelleting through a sucrose cushion. The crude ribosomal pellet was resuspended in resuspension buffer (20 mM HEPES-KOH pH 7.5, 100 mM KOAc, 5 mM Mg(OAc)_2_, 1 mM DTT, 1 mM PMSF) and treated with micrococcal S7 nuclease (Thermo Fisher Scientific, cat# EN0181) at a final concentration 40 U/mL in presence of 0.8 mM CaCl_2_ at 25°C for 15 min. The reaction was stopped by adding 2 mM EGTA. To purify 80S ribosomes, the S7-treated ribosomes were separated using 10-50% sucrose density gradient centrifugation with a SW32 Ti rotor at 125,000 *g*, 4°C for 3 h. The 80S fraction was further centrifuged in a TLA110 rotor at 335,000 *g*, 4°C for 45 min. The resulting ribosomal pellet was resuspended in 250 μL resuspension buffer (~65 A_260_/ml).

1.25 pmol of purified 80S ribosomes were subjected to test *in vitro* ubiquitination reaction with 50 μM ubiquitin, 100 nM Uba1, 300 nM Ubc4, 180 nM Not4 and energy regenerating source (1 mM ATP, 10 mM creatine phosphate, 20 μg/ml creatine kinase) and buffer-UB (20 mM HEPES-KOH pH 7.4, 100 mM KOAc, 5 mM MgCl_2_, 1 mM DTT) in 25 μl at 26°C for 60 min. The reaction was analyzed by NuPAGE and Western blotting. To prepare Ub-40S for *in vitro* reconstitution, 50 pmol of purified 80S ribosomes were used for the ubiquitination reaction as described above with respectively higher concentration of Ubc4 (5 μM) and Not4 (1 μM), incubated at 28°C for 90 min, then the tubes were placed on ice to stop the reaction. To release peptides and tRNAs from ribosomes, a high salt/puromycin treatment was performed and resulting ribosomes were loaded onto a 5-20% sucrose density gradient (20 mM HEPES-KOH pH 7.4, 500 mM KCl, 2 mM MgCl_2_, 2 mM DTT) to split the ubiquitinated ribosomes into subunits. The buffer of the 40S fraction was exchanged in an Amicon Ultra-100K to storage buffer (20 mM HEPES-KOH pH 7.4, 100 mM KOAc, 2.5 mM Mg(OAc)_2_, 2 mM DTT). Ubiquitination of eS7 was confirmed by Western blotting. The final concentration of Ub-40S was 7.6 A_260_/ml in 100 μl.

Ub-40S was used in an *in vitro* deubiquitination assay, in which 1 μl Ub-40S with 1.5 μM Otu2 in storage buffer was incubated at 30 °C, for 60 min. Presence of Ub-eS7A-HA was analyzed by Western blotting using α-HA antibodies. For reconstitution of the otu2-C178S-Ub-40S complex used for cryo-EM analysis, 12 pmol Ub-40S and 5-fold molar excess of otu2-C178S were mixed and incubated at 30°C for 10 min, placed on ice and cryo-EM grids were prepared.

### Western blotting and antibodies

After NuPAGE or SDS-PAGE, Western blotting was performed with the semi-dry method. The membrane was then incubated with 5% skim milk in phosphate buffered saline containing 0.1%w/v Tween-20 (5% milk in PBS-T) for 1 h, and further incubated with horseradish peroxidase (HRP)-conjugated specific antibodies in 5% milk in PBS-T. After 3 times washing the membrane with PBS-T, proteins were detected with the AI-600 image (GE Healthcare) using SuperSignal West Dura Extended Duration Substrate (Thermo). Antibodies targeting FLAG (1:5,000; Monoclonal ANTI-FLAG M2-peroxidase, Sigma) and HA (1:5,000; anti-HA-peroxidase, high Affinity, 3F10, Roche) were used.

### Electron microscopy and image processing

Copper grids with holey carbon support film (R3/3, Quantifoil) and a 2 nm pre-coated continuous carbon layer on top were glow discharged at 2.2 × 10^-1^ mbar for 20 s. 3.5 μl of sample was applied to the grid in a Vitrobot Mark IV (FEI Company), blotted for 2 s after 45 s of incubation at 4 °C and flash frozen in liquid ethane. Data were collected on a Titan Krios at 300 keV. For the *in vitro* reconstituted Otu2-40S complex sample, 5,726 movies were collected with a pixel size of 1.059 Å/pixel and within a defocus range of 0.5–3.2 μm using a K2 Summit direct electron detector under low-dose conditions with a total dose of 45.2 e^-^/Å^2^. For the *ex vivo* Otu2-40S complex sample, 7,765 movies were collected within a defocus range of 0.5–4.0 μm with a total dose of 46 e^-^/Å^2^. Gain-corrected movie frames were motion corrected and summed with MotionCor2^46^ and Contrast Transfer Function (CTF) parameters were determined with CTFFIND4^47^ and Gctf^48^. 540,175 (*in vitro*) and 605,551 (*ex vivo*) particle images were picked by Gautomatch with 2D projections of 40S particle images (https://www2.mrc-lmb.cam.ac.uk/research/locally-developed-software/zhang-software/#gauto). After exclusion of ice and bad particles by 2D classification on Relion 3.0 and 3.1^49^, 316,588 (*in vitro*) and 289,971 (*ex vivo*) 40S particle images were obtained. Good particles were selected, 3D refined, and classified as shown in Supplementary Fig. 3: For the reconstituted sample, 40S particles were first classified for presence of extra density for the Otu2 N-terminal hairpin on the 40S foot structure. This was done using focused classification using a soft spherical mask for this region. 10 % of particles (31,646) showed clear extra density not only for the N-terminal hairpin but also for the OTU domain of Otu2 located on eS7. This population was subjected to multi-body refinement on the 40S head and 40S body part including Otu2.

For the *ex vivo* purified sample, focused classification was first performed on the extra density for the OTU domain on eS7. Here, 22 % of particles (62,896) were enriched in Otu2, but still showed weak density for its N-terminal hairpin. A second focused classification step, now on the foot/hairpin region as yielded in a class containing 6.1 % of all particles (17,766) with extra density for the hairpin. Also here, the Otu2 enriched class (62,896 particles) was subjected to multi-body refinement. The final maps were post-processed and local resolution was estimated with Relion3.0. All ps were shown low-pass filtered according to local resolution estimation with B-factor of 20 applied in Relion3.0.

### Otu2 (150-307) purification, crystallization and structure determination

A pGEX plasmid containing the His_6_-GST-HRV 3C-tag and the gene for *S. c*. Otu2 C178S was modified by iPCR^50^ to code for the Otu2 globular domain (residues 150-307). The protein was expressed in a *E. coli* Rosetta2 DE3 strain. Cells were grown in LB media supplemented with antibiotics at 37 °C until an OD_600_ of 0.4-0.6 was reached. The temperature was then lowered to 18 °C and 0.5 mM IPTG was used for induction overnight. Cells were pelleted, resuspended in lysis buffer (50 mM sodium phosphate buffer pH 8.0, 500 mM NaCl, 10% (v/v) glycerol and 20 mM imidazole pH 8.0) and lysed using a cell homogenizer. Soluble proteins were separated from the cell debris by centrifugation (20 min, 40,000 *g*). The supernatant was incubated with Ni-NTA resin for 20 min at 4 °C, washed extensively with lysis buffer and 4 column volumes of lysis buffer containing 40 mM imidazole pH 8.0. The protein was then eluted in lysis buffer containing 300 mM imidazole and dialysed overnight at 4 °C against cleavage buffer (20 mM Tris pH 8.0, 150 mM NaCl, 1 mM DTT) and HRV 3C protease was added. To separate the His_6_-GST tag from Otu2, the sample was again incubated with Ni-NTA and the unbound fraction containing only the cleaved Otu2 globular domain was collected and concentrated to 3 mL. The sample was then loaded on HiLoad 16/600 Superdex 75 pg size-exclusion column. Peak fractions were collected, concentrated to around 20 mg/mL and flash-frozen in liquid nitrogen.

Plate-like crystals of the Otu2 C178S globular domain were obtained at 20 °C in hanging drops by mixing 1 μL of protein (22 mg/mL) and 1 μL of crystallization solution (0.2 M KSCN, 0.1 M BisTris propane pH 6.5, 20% (w/v) PEG 3350 and 20% (v/v) glycerol). Crystals were flash frozen in liquid nitrogen and crystallographic data were collected at SLS (Swiss Light Source) synchrotron. X-ray data were processed with XDS software^51^ and the structure was solved with the molecular replacement method using the program Phaser (part of the PHENIX suite)^52,53^ and the structure of the OtuD3 Otu domain (PDB ID 4BOU)^42^ as the search model. Automated model building was done with the AutoBuild software in PHENIX^54^ and manually in |COOT^55^. Restrained refinement was performed with phenix.refine of the PHENIX package^52^. Accuracy of the model was assessed using MolProbity of the PHENIX package. Final statistics can be found in Table 2.

### Model building

A composite model for the Otu2 bound to the 40S subunit with monoubiquitinated eS7 was built using the rigid body docked structure of the *S.c.* 40S subunit from PDB-6TB3^1^ as a template. The part of h44 close to the N-terminal hairpin of Otu2 was manually adjusted to fit the density using COOT ^55^. The Since the 3D reconstruction was focused on the 40S body, the resolution of the head region of the 40S was considerably lower. We therefore removed proteins and rRNA belonging to the head from our model. Due to low resolution in the platform region, we also removed proteins and rRNA belonging to the 40S platform. None of the removed parts relevant in the context of Otu2 binding. The X-ray structure of the extended OTU domain (150-307) was docked as a rigid body into the extra density on eS7. A model for Otu2-bound eS7-K83-linked monoubiquitin was obtained after superimposing the structure of ubiquitin-bound *S.c.* Otu1 followed by minor adjustments to fit the (low resolution) extra density for ubiquitin.

The Otu2 N-terminus was fitted into the cryo-EM map using a model generated by AlphaFold2. The predicted N-terminal hairpin (α1-α2; res 14-69) as well as α3 (84-94) and the long α4 fitted with only minor adjustments into the respective densities. Because the resolution did not allow unambiguous placement of sidechains of Otu2, we refrained from including them in the final model, while preserving the residue register. The AlphaFold2 model also served as a template for Otu2 loop ß3-ß4 (Ser294-Gly298) that was not resolved in the X-ray structure and was manually adjusted to fit into the cryo-EM density. Validation of the final model was done using PHENIX^52^.

### Polysome analyses

Yeast cells expressing wt or mutant Otu2-3xFLAG from plasmids were cultivated in 200 ml SDC -Leu at 30°C until an OD600 of 0.6, harvested by centrifugation and frozen in liquid nitrogen. Ground yeast cell powder - dissolved in buffer-DG containing 0.1 mg/ml cycloheximide - was centrifuged in a table-top centrifuge at 4°C to remove cell debris. The obtained lysate was then loaded on 10-50% sucrose gradient (10 mM Tris-OAc pH 7.5, 70 mM NH4OAc, 4 mM Mg(OAc)_2_) in SW40 tubes and centrifuged at 200,000 *g*, 4°C for 2.5 h. Samples were analyzed using a continuous UV detector at 260 nm connected to piston fractionator (biocomp). Each fraction was mixed well with trichloroacetic acid at final concentration 10%, placed on ice for 15 min, centrifuged at 20,000 g for 15 min, then the supernatant was completely discarded. The protein pellet was dissolved in 1x basic sample buffer (120 mM Tris, 3.5% SDS, 14 % glycerol, 8 mM EDTA, 120 mM DTT, 0.01% bromophenol Blue (BPB)), and incubated at 99°C (Otu2) or 88°C (ubiquitinated eS7A) for 10 min. Soluble fractions were analyzed by SDS-PAGE and Western blotting using α-FLAG and α-HA antibodies.

### Spot assays

Yeast cells grown in YPD for 24 h were diluted at OD600 of 0.3, and a series of 10x dilutions was prepared for each sample. 2 μL each of dilution was spotted on YPD plates in absence or presence of translation elongation inhibitors (10 μg/ml anisomycin or 50 ng/ml cycloheximide), and the plates were incubated in 30°C or 37°C for 2 days.

### Affinity purification for mass spectrometry analysis

W303-1a as a non-tag control, Otu2-FTpA, and Ubp3-FTpA yeast cells were grown in 4-6 *l* YPD until a OD_600_ of 1.5 and three independent cultures were prepared as biological replicates for both Otu2 and Ubp3. Cell were harvested by centrifugation, frozen in liquid nitrogen and lyse using a mortor and a pestle. The resulting powder was dissolved in lysis buffer 7.5 (20 mM HEPES-KOH pH 7.6, 100 mM KCl, 10 mM MgCl_2_, 1 mM DTT, 0.5 mM PMSF, 0.01% NP-40, 1 pill/13 ml of cOmplete tablets EDTA-free). After centrifugation at 40,000 *g*, 4°C for 25 min, the lysate was incubated with rabbit IgG-conjugated Dynabeads M-270 epoxy (invitrogen) for 1 h at 4°C on a nutator. The beads were then washed seven times with lysis buffer and treated with S7 nuclease (40 U/ml) at 25°C for 15 min in presence of 0.8 mM CaCl_2_. The S7 reaction was stopped by adding 2 mM EGTA, and the eluate was stored as the “S7-wash” sample. After washing the beads once by lysis buffer, the beads were incubated with His-TEV protease at 4°C for 1.5 h on a rotator. The eluate was collected as “TEV” sample. Both samples were subjected to TCA precipitation, and the protein pellet was recovered in 1x sample buffer pH 6.8 (50 mM Tris-HCl pH 6.8, 2% SDS, 10% glycerol, 0.01% BPB, 25 mM DTT). After denaturation by incubation at 95°C for 10 min and centrifugation at 20,000 *g* for 10 min, soluble fractions were analyzed by Nu-PAGE gel and stained by SimplyBlue SafeStain (invitrogen).

### Sample preparation for mass spectrometry

Affinity purified samples (n=3 per group prepared individually) were transferred to a Nu-PAGE gel and run for 6 min at 200V until the gel pockets were empty. Gels were stained for 60 min using SimplyBlue Safestain and protein-containing areas were excised. To reduce proteins, gel bands were treated with 45 mM dithioerythritol in 50 mM NH4HCO3 for 30 min at 55 °C. Free sulfhydryl groups were carbamidomethylated by 2 x 15 min incubation in a solution of 100 mM iodoacetamide in 50 mM NH4HCO3 at room temperature. Prior to digestion, gel slices were minced. Digestion was performed for 8 h at 37°C using 70 ng modified porcine trypsin (Promega, Fitchburg, WI, USA). Tryptic peptides were extracted using 70% acetonitrile and dried using a SpeedVac vacuum concentrator.

### Mass spectrometry analysis

Peptides were analyzed with an Ultimate 3000 nano-liquid chromatography system (Thermo Fisher Scientific) online-coupled to a Q Exactive HF-X mass spectrometer (Thermo Fisher Scientific). Peptides were diluted in 15 μl 0.1% formic acid and injected on an Acclaim PepMap 100 trap column (nanoViper C18, 2 cm length, 100 μM ID, Thermo Scientific). Separation was performed with an analytical EasySpray column (PepMap RSLC C18, 50 cm length, 75 μm ID, Thermo Fisher Scientific) at a flow rate of 250 nl/min. 0.1% formic acid was used as solvent A and 0.1% formic acid in acetonitrile was used as solvent B. As chromatography method a 30 min gradient from 3% to 25% solvent B followed by a 5 min gradient from 25% to 40% B was used. Data dependent mass spectrometry was performed using cycles of one full MS scan (350 to 1600 m/z) at 60k resolution and up to 12 MS/MS scans at 15k resolution. Acquired MS spectra were analyzed with MaxQuant (1.6.1.0)^56^ and the *Saccharomyces cerevisiae* subset of the UniProt database. LFQ values were used for label-free quantification. All replicates (n=3 per group) were measured twice leading to six LC-MS/MS runs per group.

### Relative protein quantification and statistics

Data analysis and statistics was done using Perseus (1.5.3.2)^57^. To handle missing values, the imputation feature of Perseus was used. For statistical evaluation, a Student’s t-test including a permutation-based FDR correction was performed. Significant hits (FDR < 0.05) with log2-fold changes < −0.6 and > 0.6 were regarded as relevant. The list of differently abundant proteins and corresponding quantitative values can be found in Supplementary Table 1.

## Acknowledgement

We thank M. Kösters, C. Ungewickell and S. Rieder for excellent technical assistance; P. Tesina for cryo-grid preparation; M. Thoms for providing yeast strains and plasmids; Y. Takehara and S. Murata for communication and discussion; the crystallization facility of the Max-Planck-Institut (Conti Department) and Dr. Jérôme Basquin for the help with the X-ray data collection.

## Funding

This study was supported by JSPS Overseas Research Fellowship to K.I., a Ph.D. fellowship by Boehringer Ingelheim Fonds to R.Bu., by grants from the JSPS [18H03977,19H05281] and AMED [20gm1110010h0002] to T. I. and by grants from the DFG to R. Be. [SFB/TRR-174, BE1814/15-1, BE1814/1-1].

## Author contributions

K.I. performed all genetic and biochemical experiments (generation of yeast strains and mutants, sucrose density gradients, spot assays), generated the *in vitro* reconstituted and *ex vivo* purified cryo-EM samples, processed and interpreted cryo-EM data. N.I. determined the crystal structure of the Otu2 extended OTU domain, J.C. and R.Bu. built and refined the Otu2-40S model, Y.M. performed initial biochemical and genetic experiments, T.F. performed comparative mass spectrometry analysis, O.B. and R.Bu. collected cryo-EM data, T.I. co-initiated the project, K.I., T.B. and R.Be. designed the study and wrote the manuscript with comments from all authors.

## Supplementary Information

Supplementary Data are available for this preprint.

Supplementary Figures 1-8

Table 1-2

Supplementary Table 1-3

Correspondence and requests for materials should be addressed to Prof. Roland Beckmann or Dr. Thomas Becker.

## Conflict of interest

All authors declare that they have no conflicts of interest.

## Notes

### Competing Interest Statement

The authors have declared no competing interest.

