## Supplemental Materials for "Recognition and deubiquitination of free 40S for translational reset by Otu2"

Table of content:

Supplementary figure. 1
Supplementary figure. 2
Supplementary figure. 3
Supplementary figure. 4
Supplementary figure. 5
Supplementary figure. 6
Supplementary figure. 7
Supplementary figure. 8

Table 1 cryo-EM statistics.
Table 2 Statistics of X-ray crystal structure.

Supplementary Table 2 Yeast strains used in this study.
Supplementary Table 3 Plasmids used in this study.

[1]Department of Biochemistry, Gene Center, Feodor-Lynen-Str. 25, University of Munich, 81377 Munich, Germany. [2]Department of Physical Chemistry, Rudjer Boskovic Institute, Bijenicka cesta 54, 10000 Zagreb, Croatia. [3]Institutes of biomedical science, Shanghai Key Laboratory of Medical Epigenetics, International Co-laboratory of Medical Epigenetics and Metabolism (Ministry of Science and Technology), Fudan university, Dong'an Road 131, 200032, Shanghai, China. [4]Graduate School of Pharmaceutical Sciences, Tohoku University, Sendai 980-8578, Japan. [5]Division of RNA and gene regulation, Institute of Medical Science, The University of Tokyo, Minato-Ku 108-8639, Japan. [6]LAFUGA, Laboratory for Functional Genome Analysis, Gene Center, Feodor-Lynen-Str. 25, University of Munich, 81377 Munich, Germany. *Correspondence to:

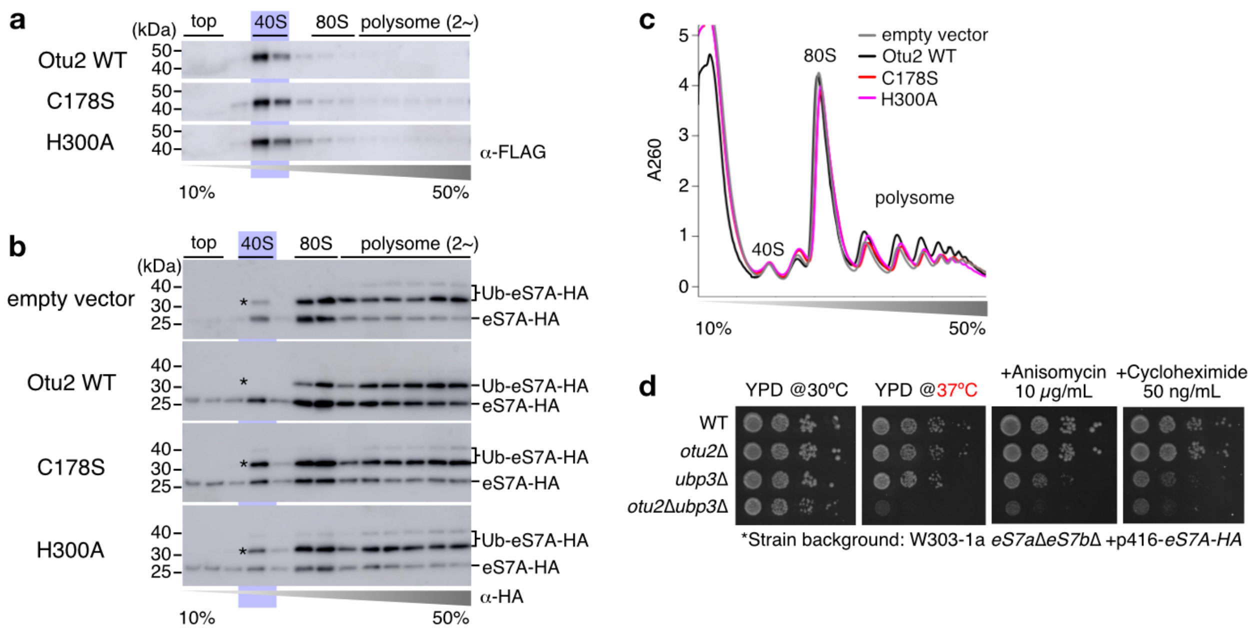

**Supplementary Fig.1 | 40S association of Otu2 and its catalytic mutants**.

a, Western Blot analysis of fractions after sucrose density gradient centrifugation. Lysates were analyzed from an *otu2*Δ with shuffled *eS7A-HA* yeast strain harboring a vector expressing wt Otu2-3xFLAG, otu2-C178S-3xFLAG and otu2-H300A-3xFLAG, respectively. Western Blots were done using α-FLAG. b, Western Blot analysis of fractions after sucrose density gradient centrifugation. Lysates were analyzed from an *otu2*Δ*ubp3*Δ with shuffled *eS7A-HA* yeast strain harboring an empty vector, a vector expressing wt Otu2, otu2-C178S and otu2-H300A, respectively. Western Blots were done using α-HA. c, UV profiles after sucrose density gradient centrifugation. d, Spot assay of yeast mutant cells in a series of 10x dilution grown for 2 days under the stress conditions.

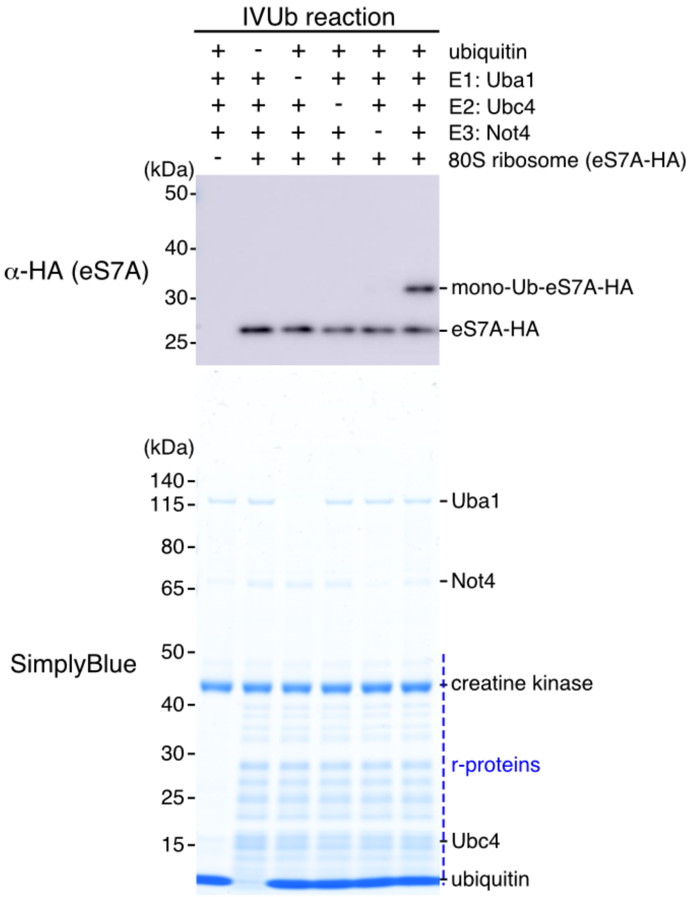

**Supplementary Fig.2 | *In vitro* monoubiquitination of eS7 in 80S ribosomes**

α-HA Western Blot monitoring *in vitro* generation of eS7A-monoubiquitinated 80S ribosomes. Below, a SimplyBlue-stained Nu-PAGE gel of the input reactions is shown.

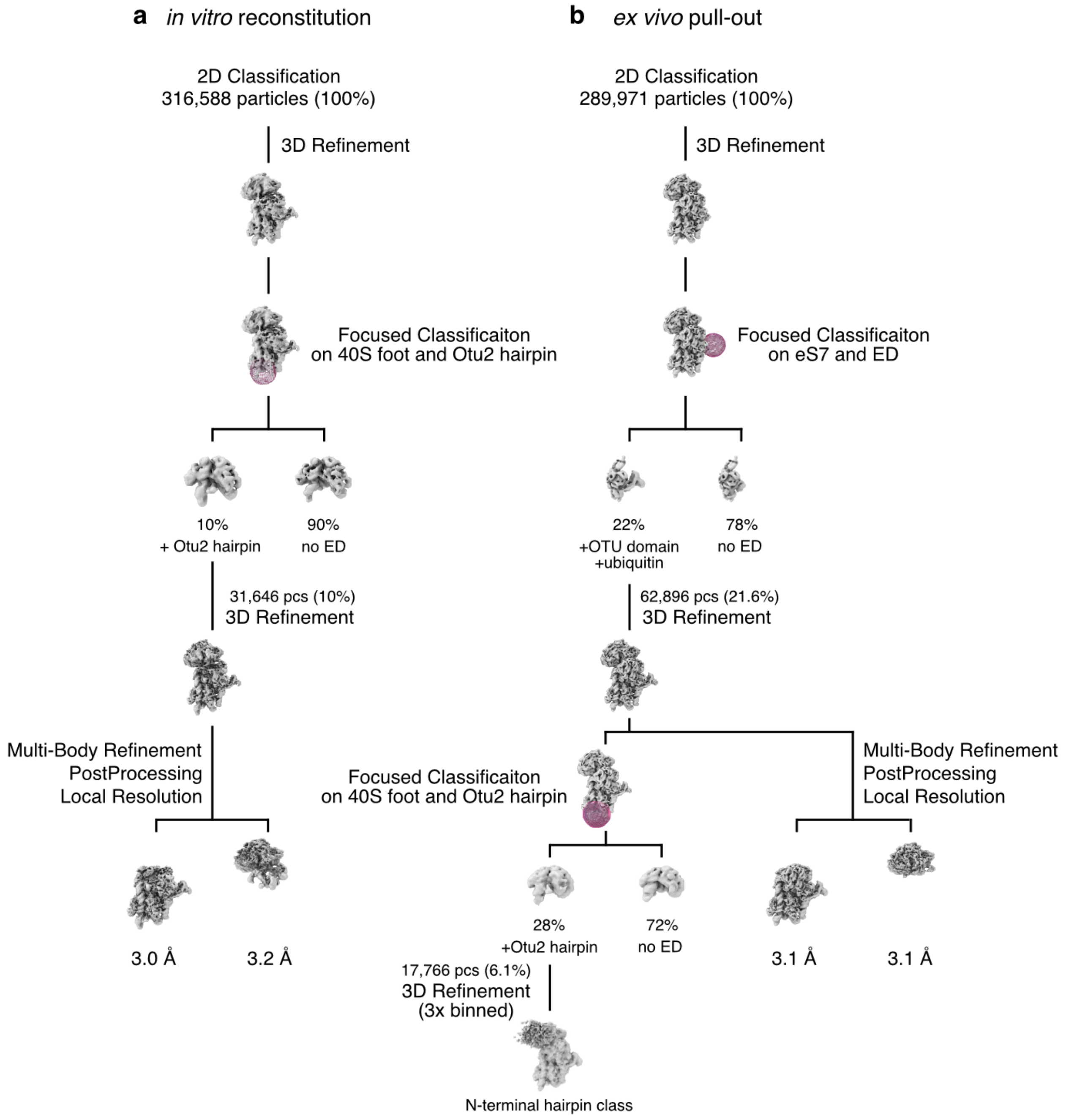

**Supplementary Fig. 3 | Data processing scheme for cryo-EM analysis of Otu2-40S complexes**

Both *in vitro* (a) and *ex vivo* (b) Otu2-40S complex datasets were processed with RELION-3.1[41]. Focused classifications using a soft spherical mask were performed on eS7 region showing extra density (ED) for the Otu2 OTU domain and ubiquitin. Further classification in the 40S foot region showed extra density for the Otu2 N-terminal domain in both samples. Classes with enriched Otu2 density were refined followed by multi-body refinement for the 40S head and 40S body region (including Otu2). This resulted in final reconstructions at an average resolution of 3.1 Å for the Otu2-containing 40S body. See Materials and Methods for details.

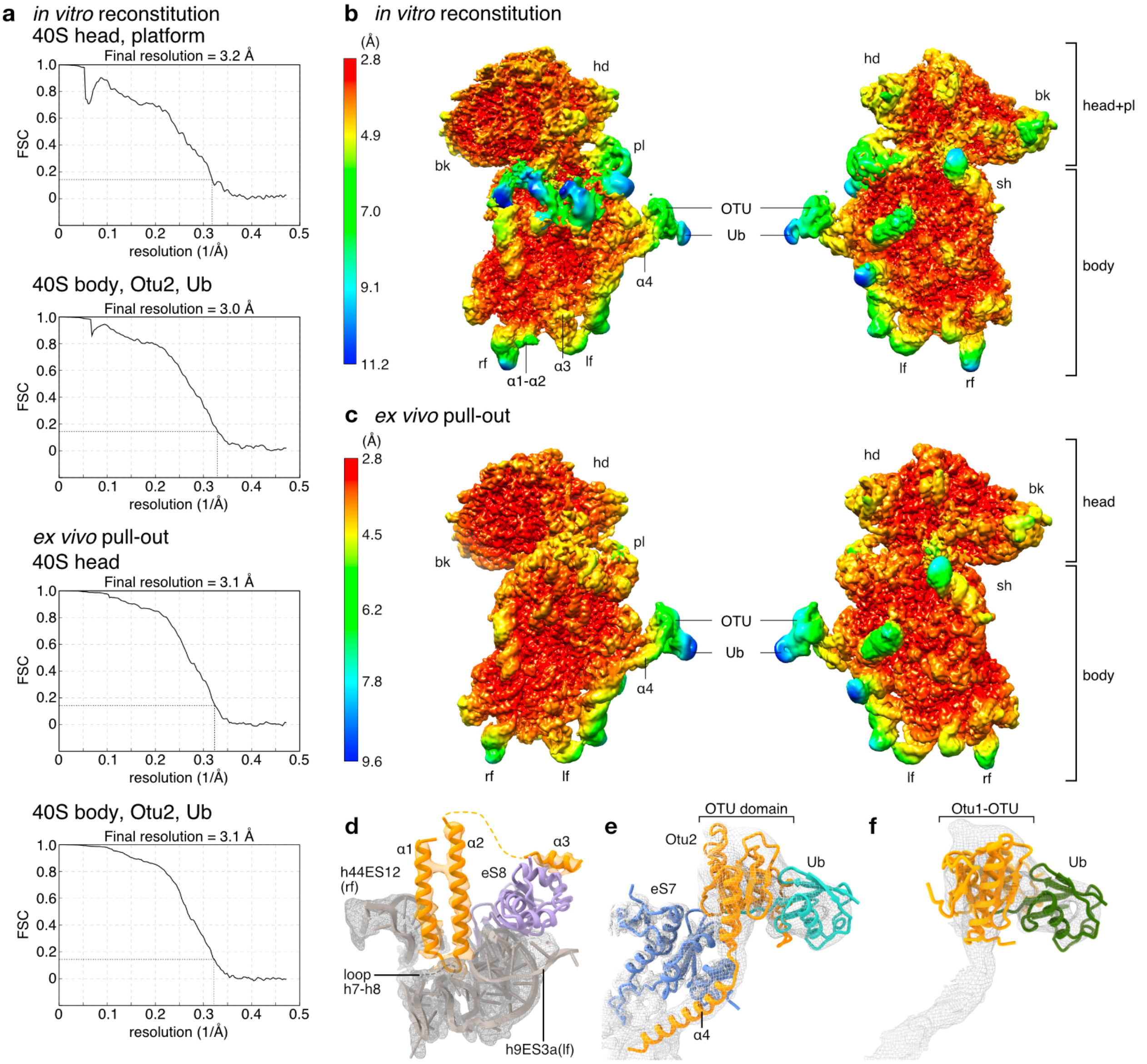

**Supplementary Fig. 4 | Resolution of Otu2-40S cryo-EM maps.**

a, Fourier shell correlation (FSC) curves for individual bodies after multi-body refinement. The average resolution was calculated according to the gold standard criterion at FSC = 0.143. b and c, Composite maps for reconstituted (b) and *ex vivo* (c) Otu2-40S complexes after multi-body refinement, colored and low-pass filtered according to local resolution estimation by RELION-3.1[41]. d-e, Models for the α1-α2 hairpin binding region (d) and α4 connecting to the OTU domain bound to Ub-eS7 fit in the respective densities (transparent mesh). f, Crystal structure of the ubiquitin-bound Otu1(3c0r;[42]) fitted into the Otu2-40S cryo-EM map. The EM maps were low-pass filtered according to local resolution.

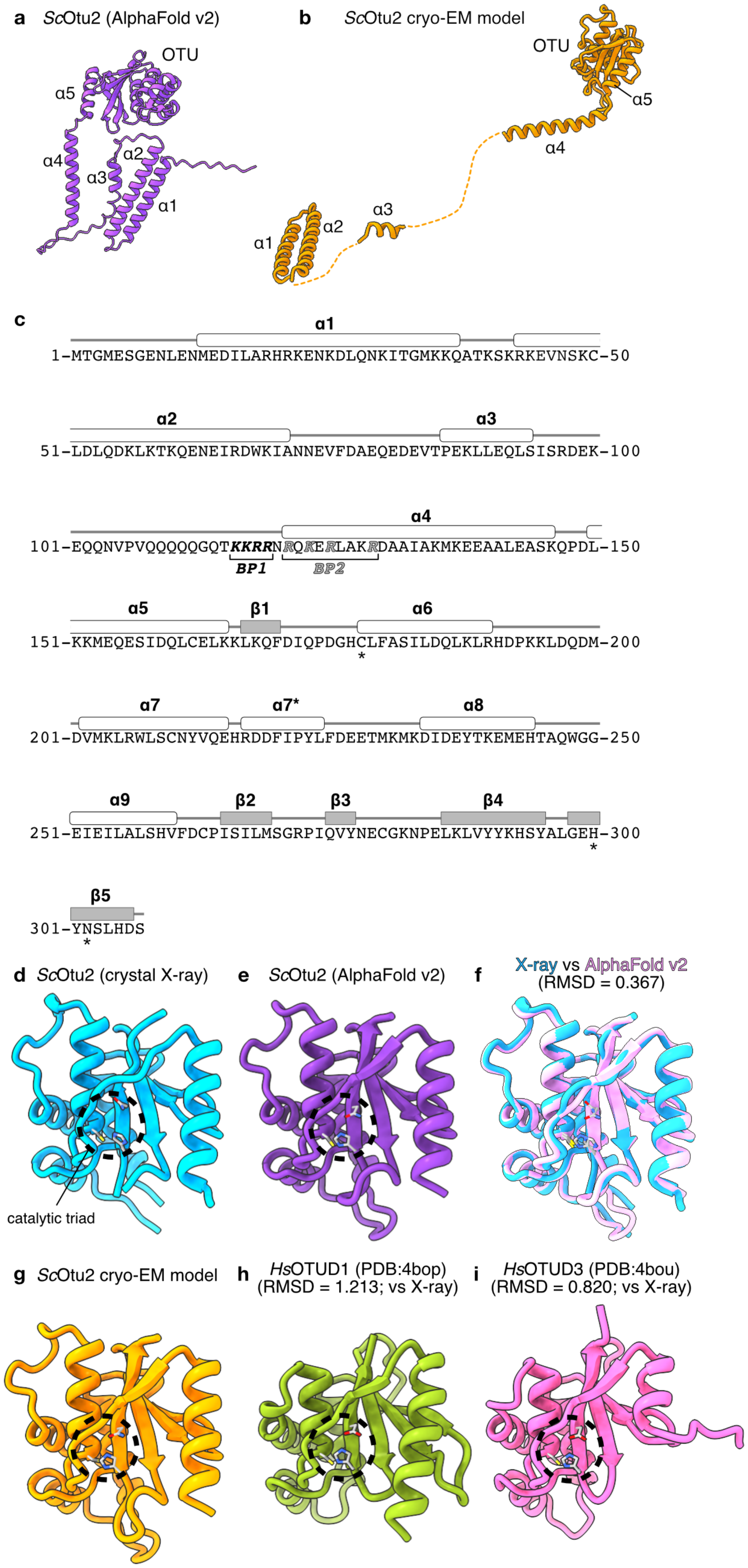

a ScOtu2 (AlphaFold v2)
OTU
α5
α4
α3
α2
α1
b ScOtu2 cryo-EM model
OTU
α5
α4
α3
α2
α1
c
α1
1-MTGMESGENLENMEDILARHRKENKDLQNKITGMKKQATKSKRKEVNSKC-50
α2
α3
51-LDLQDKLKTKQENEIRDWKIANNEVFDAEQEDEVTPEKLLEQLSISRDEK-100
α4
101-EQQNVPVQQQQQGQTKKRRNRQKERLAKRDAAIAKMKEEAALEASKQPDL-150
BP1
BP2
α5
β1
α6
151-KKMEQESIDQLCELKKLKQFDIQPDGHCLFASILDQLKLRHDPKKLDQDM-200
*
α7
α7*
α8
201-DVMKLRWLSCNYVQEHRDDFIPYLFDEETMKMKDIDEYTKEMEHTAQWGG-250
α9
β2
β3
β4
251-EIEILALSHVFDCPISILMSGRPIQVYNECGKNPELKLVYYKHSYALGEH-300
*
β5
301-YNSLHDS
*
d ScOtu2 (crystal X-ray)
catalytic triad
e ScOtu2 (AlphaFold v2)
f X-ray vs AlphaFold v2
(RMSD = 0.367)
g ScOtu2 cryo-EM model
h HsOTUD1 (PDB:4bop)
(RMSD = 1.213; vs X-ray)
i HsOTUD3 (PDB:4bou)
(RMSD = 0.820; vs X-ray)

**Supplementary Fig. 5 | Prediction of secondary structure and structural comparison for Otu2.** a, A model of full length Otu2 predicted by AlphaFold v2.0 (AF2)[4]. α1, α2, α4 and extended OTU domain showed higher per-residue confidence score (pLDDT > 70), while α3 showed low confidence (pLDDT < 70). c. b, An isolated structure of 40S-bound Otu2 determined by cryo-EM. N-terminal 4 helices widely distributed on 40S ribosome by using structurally invisible 2 linkers. c, Schematic summary of secondary structure prediction by AF2. α-helices are labeled according to their occurrence in the cryo-EM structures. The catalytic triads are indicated by asterisks. d-i, Structures of extended OTU domains. The dotted circles show the catalytic triad of deubiquitinating enzymes. d, *S. c.* Otu2 crystal structure from this study. e, *S. c.* Otu2 structure predicted by AF2. f, Comparison between the *S. c.* Otu2 crystal structure and AF2 model. g, *S. c.* Otu2 Cryo-EM model from this study. h, *H. s.* OTUD1 (PDB:4bop). i, *H. s.* OTUD3 (PDB:4bou)[41]. RMSD values in (h) and (i) refer to a comparison with the Otu2 crystal structure.

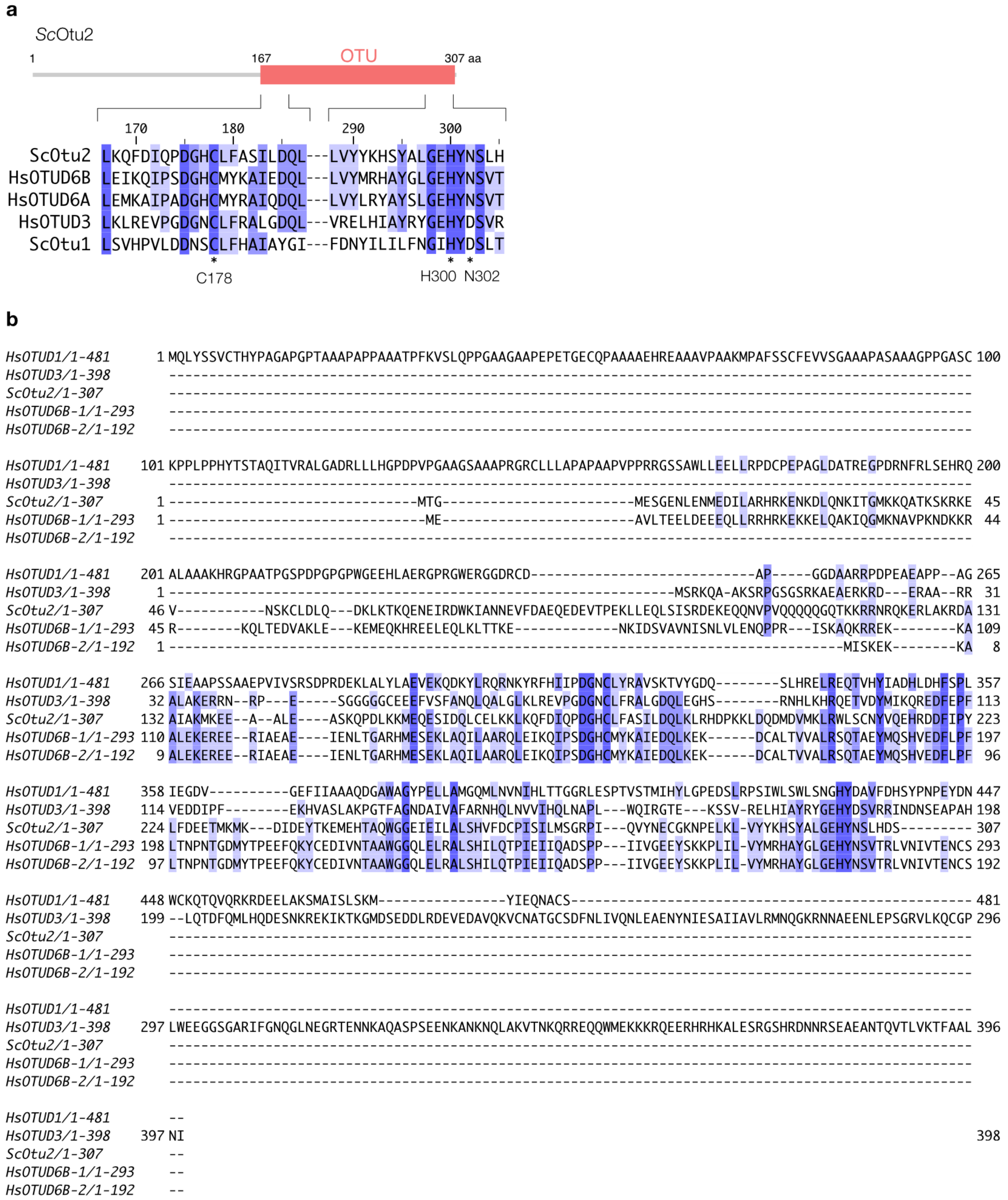

**Supplementary Fig. 6 | Sequence alignment of OTU domain containing proteins.**

a, Sequence alignment of the regions flanking the catalytic triad residues. b, Complete sequence comparison of OTU domain containing proteins. Like *H. s.* OTUD1 and OTUD3, *S.c.* Otu2 has an N-terminal S1’α-helix (Asp149-Lys165).

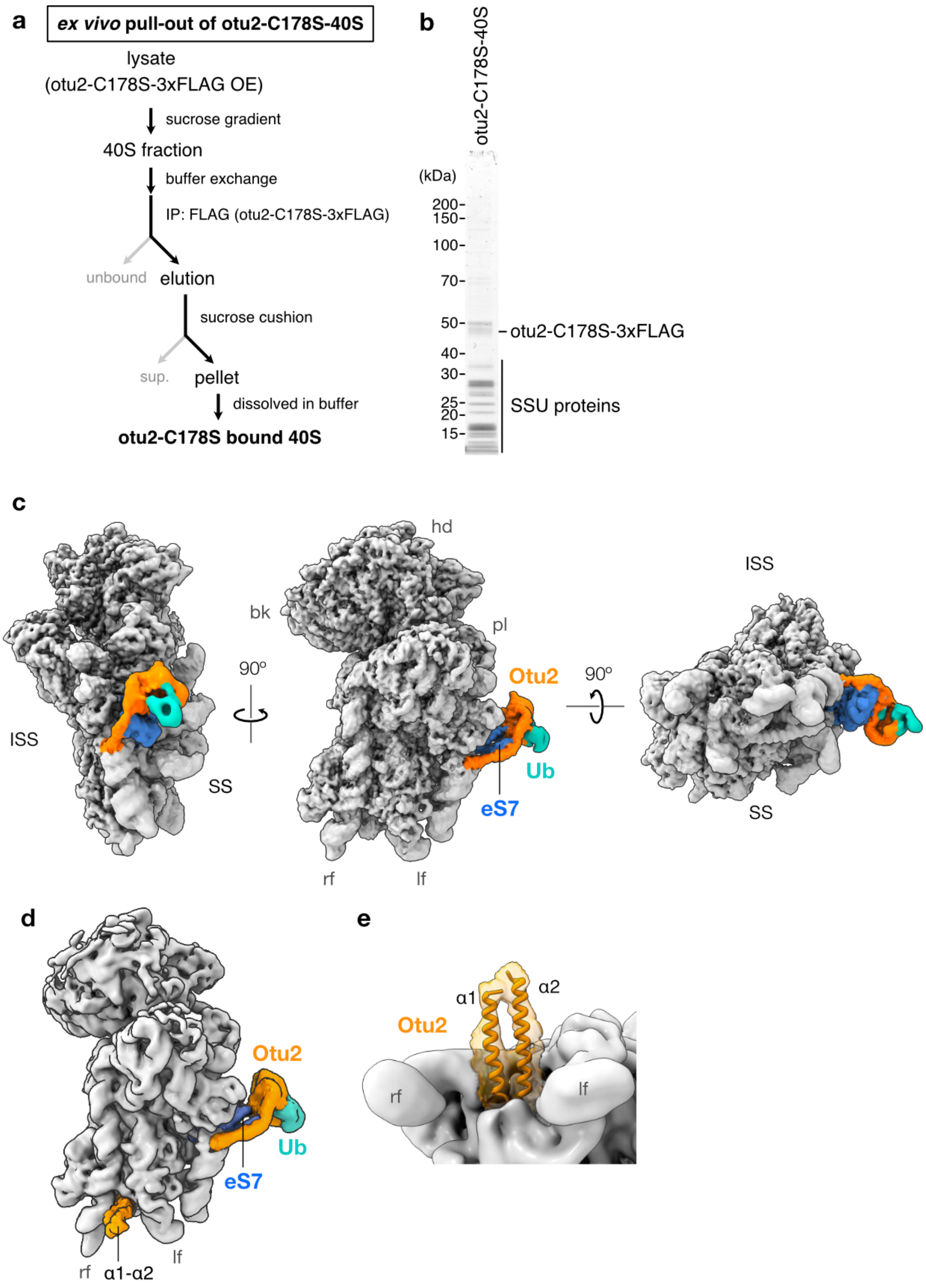

**Supplementary Fig. 7 | *ex vivo* pull-out of Otu2-40S.**

a, Scheme outlining the otu2-C178S *ex vivo* pull-out from yeast cells overexpressing otu2-C178S-3xFLAG. b, Nu-PAGE gel of the final eluate from *ex vivo* purification of otu2-C178S. Note that exclusively 40S subunit proteins co-eluted. c, Cryo-EM structure of the *ex vivo* purified otu2-C178S-40S complex. Non-ribosomal extra densities for the Otu2 OTU domain (orange) and ubiquitin (light sea green) are visible close to eS7 (blue). d, e, A subclass obtained after focused sorting on the foot region (see Supplementary Fig. 3b) showed extra density for the Otu2 α1-α2 hairpin as observed in the *in vitro* reconstituted complex.

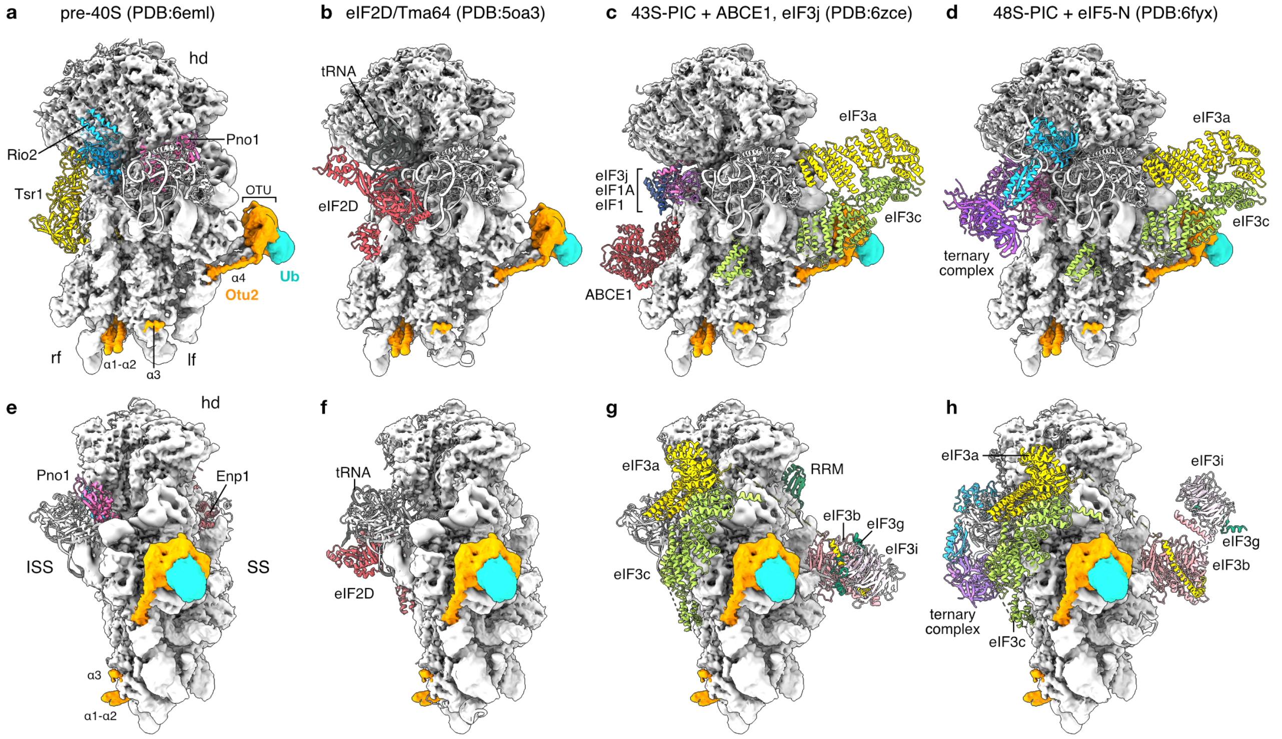

**Supplementary Fig. 8 | Comparison of 40S recycling, initiation and pre-40S complex with Otu2-40S structure.** a-h, The Otu2-40S maps (reconstituted) superimposed into models of yeast pre-40S complex (a, e; PDB: 6eml;[36]), human eIF2D-40S complex (b, f; PDB: 5oa3;[44]), yeast 43S-pre-initiation complex (PIC) with ABCE1 and eIF3j (c, g; PDB: 6zce;[45]) and yeast 48S-PIC with the eIF5 N-terminus (d, h; PDB: 6fyx;[46]).

Table 1. cryo-EM statistics

| otu2-C178S-Ub-40S | EMD-XXXX | | | |
|---|---|---|---|---|
| **Data collection and Processing** | *in vitro* reconstitution | | *ex-vivo* pull-out | |
| Magnification | 130,000 | | | |
| Voltage (kV) | 300 | | | |
| Electron exposure ($e^-/Å^2$) | 46 | | 45.2 | |
| Defocus range (μm) | 0.5 to 4.0 | | 0.5 to 3.2 | |
| Pixel size (Å) | 1.059 | | | |
| Symmetry imposed | C1 | | | |
| Micrographs Used (no.) | 5,726 | | 7,765 | |
| Initial Particle Images (no.) | 540,175 | | 606,551 | |
| Particle Images Used (no.) | 316,588 | | 289,971 | |
| Final Particle Images (no.) | 31,646 | | 62,896 | |
| Map resolution (Å) | 3.0 (body) | 3.2 (head) | 3.1 (body) | 3.1 (head) |
| FSC threshold | 0.143 | | | |
| Map sharpening *b*-factor ($Å^2$) | 20 | | | |
| **Refinement and Validation** | **PDB zzzz** | | | |
| Initial Models Used | 6TB3 (40S), 3c0r (Ub),<br>Crystal Structure and AlphaFold v2 Prediction (Otu2) | | | |
| Model Resolution (Å) | 3.7 | | | |
| FSC threshold | 0.5 | | | |
| Model Composition | | | | |
| Nonhydrogen Atoms | 45,612 | | | |
| Protein Residues | 2,725 | | | |
| Nucleotides | 1,178 | | | |
| Ligands | - | | | |
| R.m.s. deviations | | | | |
| Bond Length (Å) | 0.006 | | | |
| Bond Angles (°) | 1.02 | | | |
| Map vs. model CC (mask) | 0.72 | | | |
| MolProbity Score | 1.52 | | | |
| Clashscore | 3.17 | | | |
| Poor Rotamers (%) | 0.4 | | | |
| Ramachandran plot | | | | |
| Favored (%) | 93.89 | | | |
| Allowed (%) | 6.07 | | | |
| Disallowed (%) | 0.04 | | | |

| Table 2. Statistics for X-ray crystal structure | |
|---|---|
| **ScOtu2 extended-OTU domain C178S** | **Otu2 (150-307) C178S PDB 7PL7** |
| **Data collection** | |
| Wavelength (Å) | 0.979 |
| Space Group | I 21 21 21 |
| Cell dimension | |
| $a, b, c$ (Å) | 44.67, 60.05, 148.35 |
| α, β, γ (º) | 90, 90, 90 |
| Resolution (Å) | 42.77-2.60 (2.75-2.60) |
| Total Reflections | 86 288 |
| Unique Reflections | 11 976 |
| $R_{merge}$ | 0.139 (1.038) |
| $I/\sigma I$ | 15.1 (3.1) |
| Completeness (%) | 99.7 (98.3) |
| Redundancy | 13.0 (13.6) |
| CC1/2 | 0.999 (0.937) |
| **Refinement** | |
| $R_{work}/R_{free}$ | 0.2252/0.2643 |
| No. Atoms | |
| Protein | 1295 |
| Ligands/Ions | 0 |
| Water | 27 |
| $B$-factors | |
| Protein | 74.817 |
| Ligands/Ions | 0 |
| Water | 67.581 |
| R.m.s. deviations | |
| Bond Length (Å) | 0.003 |
| Bond Angles (º) | 0.52 |
| **Validation** | |
| Ramachandran plot | |
| Favored (%) | 93.29 |
| Allowed (%) | 6.04 |
| Disallowed (%) or Outliers (&) | 0.67 |

| Supplementary Table 2. Yeast Strains used in this study. | | | |
|---|---|---|---|
| Strain ID | Name | Genotype | Reference |
| 1 | *W303-1a wt* | *MAT* a *ade2 his3 leu2 trp1 ura3 can1* | Lab stock, parental strain |
| YKI2086 | *OTU2-FTpA* | *OTU2-FTpA-natNT2* | this study |
| YKI2157 | *UBP3-FTpA* | *UBP3-FTpA-natNT2* | this study |
| YKI2105 | *otu2* Δ | *otu2* Δ*::kanMX6* | this study |
| YKI2161 | *leu1* Δ | *leu1* Δ*::kanMX4* | this study |
| Y124 | *eS7A-HA* -shuffled | *rps7a* Δ*::HIS3MX6* , *rps7b* Δ*::natNT2* , pRS*316-RPS7A-HA-CYC1t* | Ikeuchi *et al*. 2019 |
| Y124 *otu2* Δ | *eS7A-HA* -shuffled *otu2* Δ | Y124, *otu2* Δ*::hphMX4* | this study (T. Inada) |
| Y124 *ubp3* Δ | *eS7A-HA* -shuffled *ubp3* Δ | Y124, *ubp3* Δ*::kanMX6* | this study (T. Inada) |
| Y124 *otu2* Δ*ubp3* Δ | *eS7A-HA* -shuffled *otu2* Δ*ubp3* Δ | Y124, *ubp3* Δ*::kanMX6* , *otu2* Δ*::hphMX4* | this study (T. Inada) |

| Supplementary Table 3. Plasmids used in this study. | | | |
|---|---|---|---|
| Plamid ID | Name | Expressed in | Reference/Note |
| pKI1002 | p416*GPDp-UBA1-FLAG-CYC1t* | YKI2161; *leu1* Δ | this study |
| pGEX-UBC4 | pGEX6P-1-*GST-UBC4* | *E. coli* Rossetta 2 | Ikeuchi *et al*. 2019 |
| pKI318 | pGEX6P-2-*NOT4-FLAG* | *E. coli* Rossetta 2 | this study |
| pKI1026 | pGEX6P-1-*His6-GST-3C-OTU2 wt* | *E. coli* Rossetta 2 | this study |
| pKI1027 | pGEX6P-1-*His6-GST-3C-otu2-C178S* | *E. coli* Rossetta 2 | this study |
| pOTU-Xtal | pGEX6P-1-*His6-GST-3C-otu2-C178S (150-307)* | *E. coli* Rossetta 2 | this study |
| p415GPD | p415GPD (empty vector) | Y124 *otu2* Δ, Y124 *otu2* Δ*ubp3* Δ | Lab stock T. Inada |
| pKI1034 | p415*GPDp-otu2-C178S-3xFLAG-CYC1t* | YKI2105; *otu2* Δ | this study |
| pKI1079 | p415*OTU2p-OTU2 wt-3xFLAG-CYC1t* | Y124 *otu2* Δ, Y124 *otu2* Δ*ubp3* Δ | this study |
| pKI1082 | p415*OTU2p-otu2-C178S-3xFLAG-CYC1t* | Y124 *otu2* Δ, Y124 *otu2* Δ*ubp3* Δ | this study |
| pKI1151 | p415*OTU2p-otu2-H300A-3xFLAG-CYC1t* | Y124 *otu2* Δ, Y124 *otu2* Δ*ubp3* Δ | this study |
| pKI1083 | p415*OTU2p-otu2-(1-2, 71-307)-3xFLAG-CYC1t* | Y124 *otu2* Δ*ubp3* Δ | this study |
| pKI1117 | p415*OTU2p-otu2-(1-2, 150-307)-3xFLAG-CYC1t* | Y124 *otu2* Δ*ubp3* Δ | this study |
| pKI1119 | p415*OTU2p-otu2-(1-149)-3xFLAG-CYC1t* | Y124 *otu2* Δ*ubp3* Δ | this study |
| pKI1120 | p415*OTU2p-otu2-(1-114)-3xFLAG-CYC1t* | Y124 *otu2* Δ*ubp3* Δ | this study |
| pKI1154 | p415*OTU2p-otu2-K116A K117A R118A R119A-3xFLAG-CYC1t* | Y124 *otu2* Δ*ubp3* Δ | this study, BP1-mut |
| pKI1155 | p415*OTU2p-otu2-R121A K123A R125A R129A-3xFLAG-CYC1t* | Y124 *otu2* Δ*ubp3* Δ | this study, BP2-mut |
